# HOGVAX: Exploiting Peptide Overlaps to Maximize Population Coverage in Vaccine Design with Application to SARS-CoV-2

**DOI:** 10.1101/2023.01.09.523288

**Authors:** Sara C. Schulte, Alexander T. Dilthey, Gunnar W. Klau

**Affiliations:** Algorithmic Bioinformatics, Heinrich Heine University Düsseldorf, Germany; Institute of Medical Microbiology and Hospital Hygiene, University Clinic Düsseldorf, Germany

**Keywords:** peptide vaccine design, SARS-CoV-2, combinatorial optimization, hierarchical overlap graph, string problem

## Abstract

Peptide vaccines present a safe and cost-efficient alternative to traditional vaccines. Their efficacy depends on the peptides included in the vaccine and the ability of major histocompatibility complex (MHC) molecules to bind and present these peptides. Due to the high diversity of MHC alleles, their diverging peptide binding specificities, and physical constraints on the maximum length of peptide vaccine constructs, choosing a set of peptides that effectively achieve immunization across a large proportion of the population is challenging.

Here, we present HOGVAX, a combinatorial optimization approach to select peptides that maximize population coverage. The key idea behind HOGVAX is to exploit overlaps between peptide sequences to include a large number of peptides in limited space and thereby also cover rare MHC alleles. We formalize the vaccine design task as a theoretical problem, which we call the Maximum Scoring k-Superstring Problem (MSKS). We show that MSKS is NP-hard, reformulate it into a graph problem using the hierarchical overlap graph (HOG), and present a haplotype-aware variant of MSKS to take linkage disequilibrium between MHC loci into account. We give an integer linear programming formulation for the graph problem and provide an open source implementation.

We demonstrate on a SARS-CoV-2 case study that HOGVAX-designed vaccine formulations contain significantly more peptides than vaccine sequences built from concatenated peptides. We predict over 98% population coverage and high numbers of per-individual presented peptides, leading to robust immunity against new pathogens or viral variants.

## 1 Introduction

Vaccination provides an effective way to develop immunity against many diseases. These include the infectious respiratory tract disease COVID-19, which is caused by variants of the coronavirus SARS-CoV-2. Fortunately, many vaccines already exist, but further research is needed to be able to quickly react in the face of viral mutations and to develop better vaccines against current and upcoming diseases. In addition, the efficacy of vaccines can be improved by targeting immunity in larger fractions of the population. This is difficult because the genetic makeup of the human immune system is highly diverse.

Modern vaccination approaches like mRNA or peptide vaccines allow for targeted immunization with fewer side effects than traditional vaccines by live attenuated or inactivated pathogens [4,14]. Targeting allows to focus only on those parts of a pathogen that trigger immune responses, whereas other parts might contribute only little to the immunizing effect.

While mRNA vaccines contain messenger RNA of whole viral proteins, the components of peptide vaccines are small, synthetically produced fragments from proteins, i.e., peptides, of a target pathogen that are known for being immunogenic. This allows for an even more tailored approach and an increased population coverage. Peptides of the nucleocapsid (N) protein of SARS-CoV-2 are, for example, promising candidates for vaccine design [18], while presenting the whole N protein can cause inflammatory lung injuries by interacting with a signaling pathway of the immune system [8]. In this work, we focus on the design of peptide vaccines.

The efficacy of a peptide vaccine essentially depends on two components: the peptides included in the vaccine and the ability of major histocompatibility complex (MHC) molecules to bind and present these peptides to cells of the immune system. Due to the high diversity of MHC alleles and their diverging specificities in binding peptides, choosing a set of peptides that maximizes population coverage is a challenging task. Further, peptide vaccines are limited in their size allowing only for a small set of peptides to be included. Thus, they might fail to immunize a large part of the human population or protect against upcoming viral variants [19].

Bioinformatics has long been well established in vaccine development. Toussaint et al. presented an approach to select a diverse set of peptides that maximize immunogenicity [20]. Vider-Shalit et al. compute a peptide vaccine sequence that maximizes population coverage and induces a large number of peptides presented by human leukocyte antigen (HLA) alleles, i.e., peptide-HLA hits, by further optimizing the peptide cleavage probability for each chosen peptide [21]. Schubert and Kohlbacher follow the approach to design peptide vaccines by concatenating peptides with spacer sequences in between [17]. This is called a string-of-beads vaccine, where the beads should ensure that the peptides are correctly cut by the proteasome [17]. Recently, Liu et al. [13] developed OptiVax, a heuristic method to choose a set of peptides that maximizes population coverage based on the HLA allele frequencies of the target population.

In this contribution, we build on the work by Liu et al. [13] and address the major challenge of selecting an optimal set of peptides to be included in the vaccine that maximizes population coverage, i.e., the fraction of individuals in a population that is immunized by the vaccine. Our key idea is to exploit overlaps between peptide sequences to include a large number of peptides in a limited space and thereby also cover rare MHC alleles. We model this task as a theoretical problem, which we call the *Maximum Scoring k-Superstring Problem* (MSKS). We show that MSKS is NP-hard and introduce a reformulation as a graph problem using the hierarchical overlap graph (HOG) data structure. We give an integer linear programming (ILP) formulation for the graph problem. Using the SARS-CoV-2 case study from [13] we show that our vaccine formulations contain significantly more peptides compared to vaccine sequences built from concatenated peptides, including the state-of-the-art tool OptiVax. We predicted over 98 % population coverage for our vaccine candidates of MHC class I and II based on single-allele and haplotype frequencies. Moreover, we predicted high numbers of per-individual presented peptides leading to robust immunity in the face of new virus variants. HOGVAX, our novel approach, is publicly available as an open source implementation.

## 2 MSKS: Combinatorial Model for Maximum Population Coverage Vaccine Design

We now introduce our novel combinatorial model for peptide vaccine design, which requires three types of input: (i) candidate peptide sequences, (ii) HLA allele frequencies, and (iii) binding affinity predictions between peptides and HLA alleles. To obtain the input data for the SARS-CoV-2 case study we follow the procedure by Liu et al., which is described in detail in Section 4.1.

1. The peptide candidates stem from selected regions of the pathogen proteome and are usually generated with a sliding window approach, followed by a filtering step to remove unwanted candidates such as peptides that are likely to mutate. We model candidate peptides at the sequence level, that is, we consider them as a set of short strings.
2. The distribution of HLA alleles among the target population plays a key role when designing a vaccine. The frequencies of HLA alleles are available from databases such as dbMHC [12], as obtained from the IEDB Population Coverage Tool download [3]. In addition, linkage between the alleles can be considered by combining the frequencies of allele combinations with haplotype or genotype data of typed individuals. Here, we assume a frequency vector for each allele combination under consideration.
3. The immunizing effect of a peptide vaccine that induces T cell activation depends upon MHC class I and II antigen presentation. Thus, it is crucial to know the probability of a specific peptide being presented by an MHC molecule. As the binding affinities are difficult to obtain experimentally, we use in silico prediction methods to obtain these probabilities. We assume a representation of this information as a binary matrix expressing whether a specific candidate peptide binds to an allele combination.

Given such input, that is, a set of candidate peptides, the allele frequencies and binding affinities, the task is now to choose which of the peptides to put into the vaccine. We formulate the following two criteria such a selection should meet:

– The length of the selected peptides should fit into a given length limit *k*. Here, we exploit possible overlaps between the peptide sequences. In order to use the space most efficiently we therefore aim at finding a shortest common superstring of the selection.
– The selection should cover a large part of a given population in the sense that at least one selected peptides binds to an MHC allele.

We therefore introduce the following combinatorial problem:

### Definition 1 (Maximum-Scoring *k* Superstring Problem, MSKS)

*Given a set of strings S* = *{s*_1_, *s*_2_, …, *s*_*n*_*} over a finite alphabet ∑, a frequency vector f* = (*f*_1_ *f*_2_ … *f*_*m*_)^*T*^ *with m* ∈ *ℕ and f*_*a*_ ∈ [0, 1] *for a* = 1, …, *m, an m × n binary matrix B, and a limit k* ∈ℕ. *We are searching for P* ⊆ *S and a superstring* 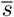 *of P, such that*

1. *the length of* 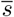 *is at most k, and*
2. *the population coverage* 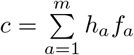 *is maximal, where*

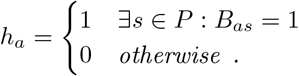

We call such a string 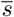 a *maximum-scoring k superstring*. The following theorem states that the problem to compute such a string is intractable.

### Theorem 1.

*The decision variant of MSKS is NP-complete*.

*Proof*. The decision variant of MSKS has an additional parameter *c*′ and the question is whether a solution exists with a common superstring of length at most *k* and with population coverage of at least *c*′.

The problem is clearly in NP as, for a given solution (*P, s*), we can easily verify in polynomial time whether *s* is a superstring of *P* of length at most *k* and whether the population coverage of *P* exceeds *c*′.

To show hardness, we give a reduction from the well-known NP-hard shortest common superstring (SCS) problem [9], whose decision is defined as follows: Given a set of strings *S* = {*s*_1_, …, *s*_*n*_} and a length *k*′, is there a common superstring 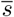 of all strings in *S* with length at most *k*′?

Given an instance of the SCS problem, we construct an instance of the decision version of MSKS as follows: We set *k* = *k*′ and *c*′ = 1. We build an *n*-dimensional frequency vector *f* with *f*_*a*_ = 1*/n* for each *a*∈ {1 …, *n*} and set *B* to the binary identity *n ×n*-matrix. All transformations take polynomial time.

If a solution for SCS with length *k* exists, a solution for the MSKS with length *k* and score 1 exists. Vice versa, if we have a solution for MSKS with score 1, we have selected all strings in *S* within the given limit of *k*. Thus, a solution for SCS with length *k* exists, too. Consequently, the SCS instance has a yes-answer if and only if the transformed MSKS instance has a yes-answer. □

## 3 HOGVAX: Solving MSKS using Hierarchical Overlap Graphs

Our approach is based on a reformulation of MSKS as a graph problem. The shortest common superstring problem is a well-studied problem and can, for example, be expressed as a shortest Hamiltonian path problem in an appropriately weighted overlap graph. Because of the quadratic number of edges in the overlap graph and the large number of peptides we consider, we decided to base our reformulation on the more space-efficient *hierarchical overlap graph* (HOG), which was first proposed by Cazaux and Rivals [5].

Instead of a fully connected overlap graph, the HOG is a directed tree-like structure that represents overlaps as inner nodes. It is a compressed version of the Aho-Corasick trie [1]. Park et al. [15] showed how to construct the HOG of a set of strings in linear time and space. Originally, the data structure is defined on substring-free input sets. In our setting, this does not apply as, for two input peptides *p*_1_ and *p*_2_ where *p*_1_ is a substring of *p*_2_ we may want to choose only *p*_1_ and not *p*_2_. We therefore extend the HOG definition and linear-time construction algorithm to allow for this generalized input.

Fig. 1 shows the Aho-Corasick trie and the HOG *H* = (*V, E*) for a small set of four peptides. Both data structures capture the input strings as paths in a tree and contain *suffix links* pointing to the longest suffix that is also a prefix of some input string. Each node *v* ∈ *V* corresponds to a prefix of an input string *L*(*v*) formed by following the edges from the path from the root to *v*. The *length* of an edge *ij* ∈ *E* is defined as

**Fig. 1.**
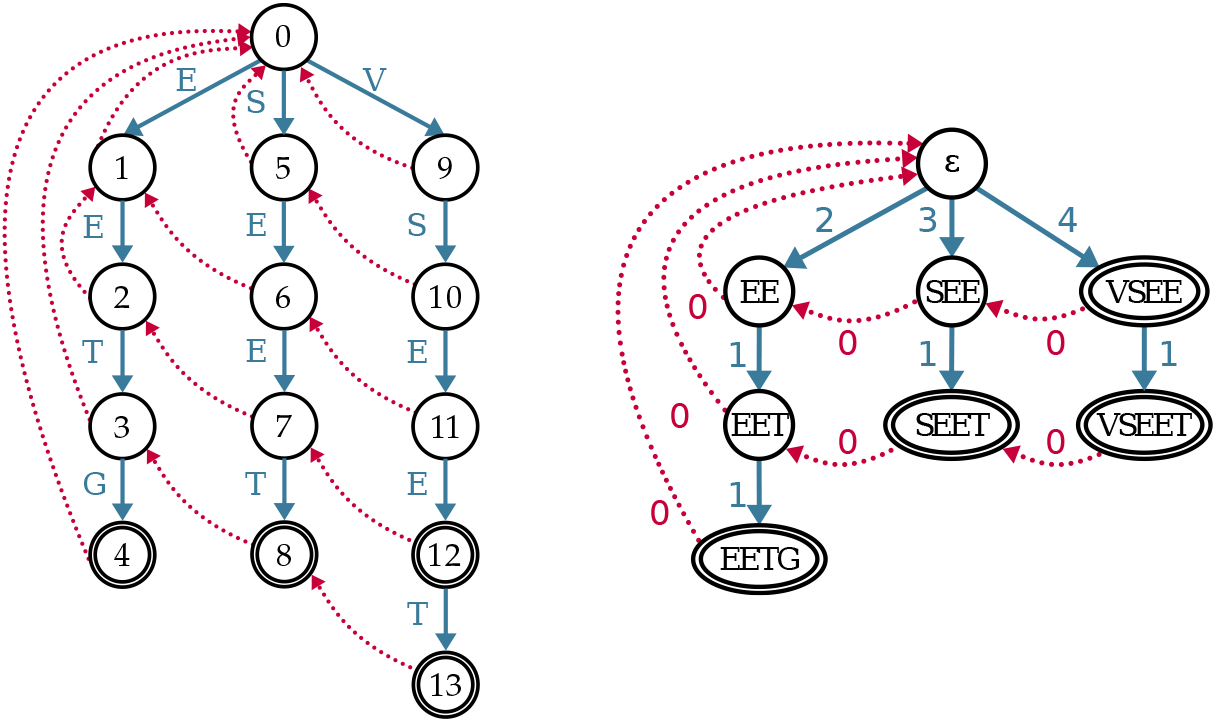
The Aho-Corasick trie (left) and HOG (right) for the string set *S* = {EETG, SEET, VSEE, VSEET} with tree edges colored in blue and suffix links in red (dashed). Double-encircled nodes correspond to whole strings of the input set.

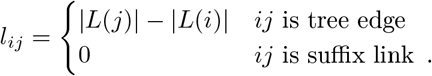

We can now reformulate MSKS using the HOG data structure. A superstring corresponds to a directed closed walk through the HOG that starts and ends at the root node. Note that a walk may contain nodes and edges repeatedly.

### Lemma 1.

*Given a set of strings S with corresponding HOG H* = (*V, E*) *rooted at r* ∈*V*. *A closed walk C through H that starts and ends at r corresponds to a superstring* 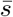 *of all the strings {L*(*v*) | *v* ∈ *C and s* ∈ *S} with* 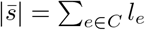.

*Proof*. The closed walk can be decomposed into collection of cycles starting and ending at the root. Following each cycle we “collect” strings *s* ∈*S* whenever we pass through a node *v* ∈*V* with *L*(*v*) = *s* by a path from *r*. The sum of the edge lengths of this path is equal to the length of *s* and the length of the cycle is equal to the length of the superstring that contains all collected input strings. Each cycle closes with a suffix link of length zero. □

### Definition 2 (Maximum-Scoring *k* Walk Problem, MSKW)

*Given an MSKS instance* (*S, B, f, k*) *and the HOG H* = (*V, E*) *for the string set S. We are searching for a closed walk C in H starting and ending at the root r* ∈ *V such that*

1. *the sum of the edge weights of C is at most k, and*
2. *the population coverage* 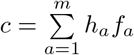 *is maximal, where*

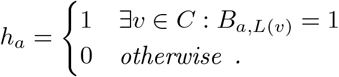

It follows immediately from Lemma 1 that MSKW is a proper reformulation of MSKS and that the proplems are, in essence, identical. Therefore, we concentrate on solving MSKW, for which we give an integer linear programming (ILP) formulation.

Each edge *ij* of the HOG *H* = (*V, E*) is assigned to an integer decision variable *x*_*ij*_, where *x*_*ij*_ gives the number of times edge *ij* is traversed. Our goal is to maximize population coverage by choosing peptides that cover the alleles of a target population. Therefore we associate each allele *a* ∈{1, …, *m*} with a binary variable *h*_*a*_. An allele is covered if at least one peptide from the vaccine set binds to the allele. By Def. 2, this is the case if the walk contains at least one node of the HOG allocated to one of the input peptides for which the entry in the binding affinity matrix *B* is 1 and therefore *h*_*a*_ = 1. Otherwise the allele is uncovered and *h*_*a*_ = 0. A node is included in the walk if an edge entering the node and an edge leaving the node are traversed by the walk. The ILP is as follows:

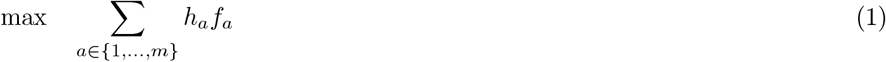

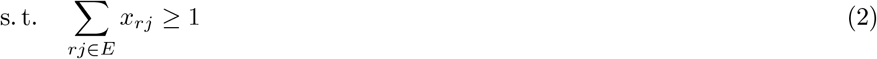

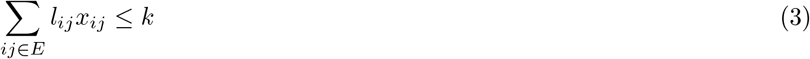

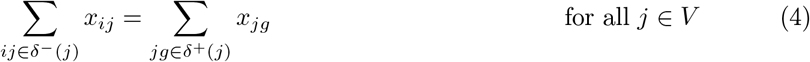

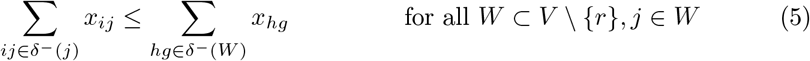

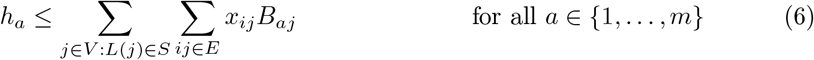

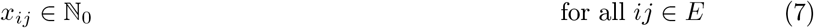

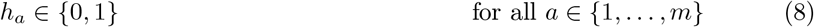

We address maximizing population coverage by maximizing the allele frequency of covered alleles and use the term given in Def. 2 as the objective function (1) of HOGVAX. As we search for a closed walk starting from the root node, we require at least one outgoing edge from the root *r* to start the walk (2). Whenever we traverse an edge *ij*, i.e., *x*_*ij*_ ≥ 1, we must add the weight *l*_*ij*_*x*_*ij*_ to the total length of the vaccine sequence constructed from the traversal. The sum of edge weights must not exceed the given limit of *k* (3). Further, to build a closed walk, we must assure that for each visit of a node, we enter the node and leave it again. Thus, we apply a flow conservation constraint for each node ensuring that the number of incoming edges equals the number of outgoing edges (4). We denote the set of incoming edges of a node *j* by *δ*^−^(*j*) and the set of edges leaving *j* by *δ*^+^(*j*).

We intend to construct a single vaccine sequence described by a connected walk. Thus, we use a subtour elimination constraint to avoid additional cycles that are not reachable from the root node. We reformulate the generalized subtour elimination constraint given in [2] as follows. For each subset of nodes *W*⊂*V\ r*, we require that for each incoming edge to a node *j* ∈*W* there must be an incoming edge to *W* (5). More precisely, if an edge *ij* for *j* ∈*W* is in the walk, then there must be an edge *hg*∈ *δ*^−^(*W*) with *x*_*hg*_ ≥1 where we define *δ*^−^(*W*) = {*hg*∈ *E* |*g* ∈*W, h* ∉ *W*}.

Finally, we define the constraints on our *h*_*a*_ variables. Consider a node *j* ∈*V* that represents the peptide sequence *L*(*j*) ∈ *S*. The node is part of the walk if an incoming edge *ij* is in the walk, thus if any *x*_*ij*_≥ 1. An allele *a* is covered, i.e., *h*_*a*_ = 1, if an *x*_*ij*_≥ 1 exists for which *B*_*aj*_ = 1 as well. We must sum over all nodes representing peptides, and in a second sum, go over all incoming edges to address each *x*_*ij*_ which is multiplied with the corresponding matrix entry *B*_*aj*_. We add such a constraint (6) for each allele of our input frequencies.

The general HOGVAX workflow is shown in Fig. 2. Note that we adapt the binding affinities to account for input peptides that are substrings of other input peptides. As this work is based on the assumption that the proteasome cuts out all candidate peptides of the vaccine sequence, we expect the substring is cut from the superstring when the larger string is selected. Therefore, we merge the scores of an allele and all peptide sequences in the binding affinity prediction matrix such that a superstring is given the maximum score of itself and its substrings.

**Fig. 2.**
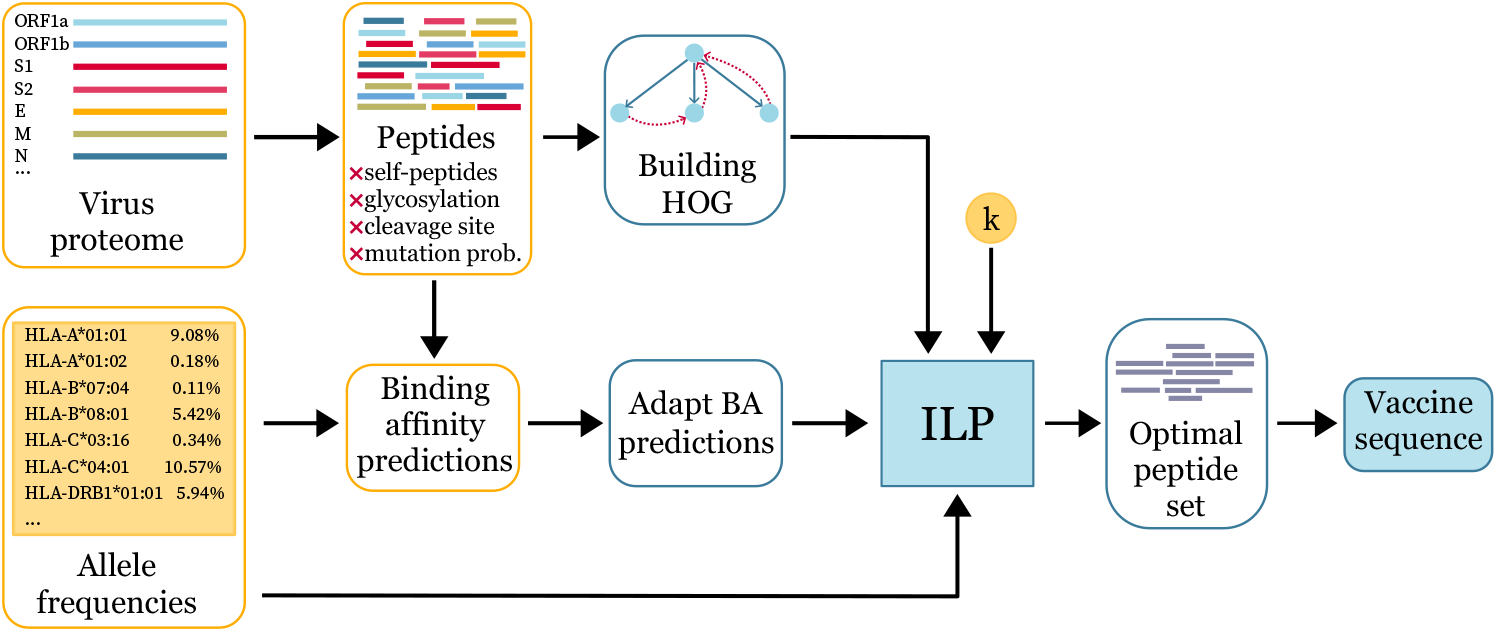
Yellow parts refer to input data, blue parts represent the HOGVAX workflow. We build a generalized HOG from the peptides, adapt the binding affinities (BA) and use this, together with the allele frequencies and the length limit *k*, to construct our ILP, the solution of which is an optimal selection of candidate peptides as well as their superstring of length at most *k*.

## 4 Experimental Results

We compare HOGVAX to OptiVax [13], the state-of-the-art tool for population coverage maximization on a case study on SARS-CoV-2, which has also been introduced in [13]. Here, we compare MHC class I and class II vaccines with respect to population coverage using different metrics. To determine *k*, the maximum length of the HOGVAX vaccine, we used the length of the concatenated output peptides from OptiVax.

The source code for HOGVAX and necessary input files, results, computed vaccine sequences, preprocessing and evaluation scripts are available for download at https://gitlab.cs.uni-duesseldorf.de/schulte/hogvax. The computations were executed on a workstation with the following specification: AMD EPYC 7742 64-Core processor with 128 threads, 1 TiB DDR4 RAM, running Debian Buster with Linux kernel 5.8.

### 4.1 SARS-CoV-2 case study data

We obtain peptide sequences, HLA allele frequencies, and binding affinity predictions between pairs of peptides and HLA alleles as described in Liu et al. [13]: The peptides were created from the SARS-CoV-2 proteome using the official reference entry Wuhan/IPBCAMS-WH-01/2019 from the GISAID database [7] that provides the nucleotide sequences of the open reading frames (ORFs) and the four viral structural proteins E, M, N, and S. Using a sliding window approach results in 29 403 MHC class I peptides and 125 593 MHC class II peptides. We filter the peptide set as in [13]. This includes the elimination of self-peptides, glycosylated peptides, peptides of cleavage regions, and peptides that are likely to mutate. After filtering we obtain a set of 4 512 MHC class I peptides and 37 435 MHC class II candidate peptides.

Allele sequences of single HLA alleles were collected from the dbMHC database [12], their frequencies were accessed with the IEDB population coverage tool [3]. The data contains frequencies for 2 395 alleles across 239 geographical regions for MHC class I, and frequencies for 275 alleles across 274 geographical regions for MHC class II. Liu et al. also considered linkage between the loci. For this, MHC class I and II alleles were determined with high-resolution typing for 2 327, 2 886, and 1 653 individuals with self-reported European, African, and Asian ancestry, respectively [13]. The data is composed of 2 138 independent MHC I haplotypes with 230 distinct HLA-A, HLA-B, and HLA-C alleles, and 1 711 MHC II haplotypes with 280 different HLA-DP, HLA-DQ, and HLA-DR alleles. All allele and haplotype frequencies are normalized to sum up to 1 for each locus.

Binding affinity predictions are calculated with NetMHCpan-4.1 and Net-MHCIIpan-4.0 for MHC class I and II, respectively [16]. Predictions were generated for each combination of 233 HLA class I alleles and 29 402 peptides of length 8 to 10, and for each combination of 283 HLA class II alleles and 125 593 peptides of length 13 to 25. As in Liu et al. we define a binding affinity threshold of 0.638, which corresponds to a concentration of 50 nM.

### 4.2 OptiVax

The method by Liu et al. [13] aims at choosing a subset of the candidate peptides that maximizes population coverage based on the HLA allele frequencies of the target population. It consists of two components: OptiVax, to optimize coverage, and EvalVax, to evaluate solutions. OptiVax uses beam search to heuristically explore the solution space. The beam search starts with an empty set that is iteratively extended by single peptides. In each iteration, only the top *b*-many solutions are explored further, where *b* is the so-called beam size. After each iteration, the current candidate sets are evaluated by EvalVax. In the first iteration, all input peptides are evaluated. Only the peptides with the *b* highest EvalVax-scores are considered further and expanded with a further peptide. The process is repeated until a minimal set with the desired population coverage is computed or the pre-defined maximum of allowed peptides is reached. Vaccine candidates for MHC classes I and II are designed separately with a beam size of *b* = 10 and *b* = 5 for MHC class I and II, respectively.

### 4.3 Evaluation metrics

We evaluated the HOGVAX and OptiVax vaccine candidates using the EvalVax evaluation functions. Note that these are the objective functions used by OptiVax. Liu et al. suggested EvalVax-Unliked for evaluating a peptide selection *P* in the single allele frequency setting with independent loci. They define and suggest to use the population coverage for peptide selection *P* as 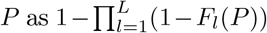 where *F*_*l*_(*P*) is the probability that an individual displays at least one peptide of the vaccine at locus *l* within a set of *L* loci in total. We refer to this measure as *unlinked coverage C*_*u*_.

Liu et al. also suggest EvalVax-Robust, an evaluation metric that takes linkage disequilibrium between alleles of distinct loci into account. In this case, they suggest to use the probability that an individual has at least *h* peptide-HLA hits. We refer to this measure as *robust coverage C*_*r*_. For details of the EvalVax metrics, see [13].

### 4.4 Implementation of HOGVAX and runtime experiments

HOGVAX is implemented in Python. We use it with the software management system Conda for an easy and reproducible execution of HOGVAX. For solving the ILP we use the Gurobi Optimizer [11] that provides a Python API. We used a separation approach for the subtour elimination constraints via a Gurobi callback. In all experiments, we run the solver until it finds an optimal variable assignment. HOGVAX computes provably optimal vaccine sequences in under 15 minutes for all instances but the MHC II haplotype instance for which HOGVAX takes about 1.3 hours. OptiVax finishes all computations on our workstation in under 25 minutes.

### 4.5 Comparison of population coverage on single allele frequencies

For the comparison of single-allele MHC class I vaccines, we use allele frequencies of an average world population. We first ran OptiVax-Unlinked, resulting in a vaccine candidate of 19 peptides. From their concatenation, we inferred *k* = 170 as limit for the vaccine candidate computed with HOGVAX. The HOGVAX MHC class I solution contained 97 peptide candidates from seven distinct SARS-CoV-2 proteins, which was about five times the number of the OptiVax peptides, see Fig. 3A. In contrast, OptiVax-Unlinked selected peptides from four different proteins. In both methods, peptides from the ORF1a and ORF1b proteins dominated the sets. The exact compositions are shown in Fig. 3C and D.

**Fig. 3.**
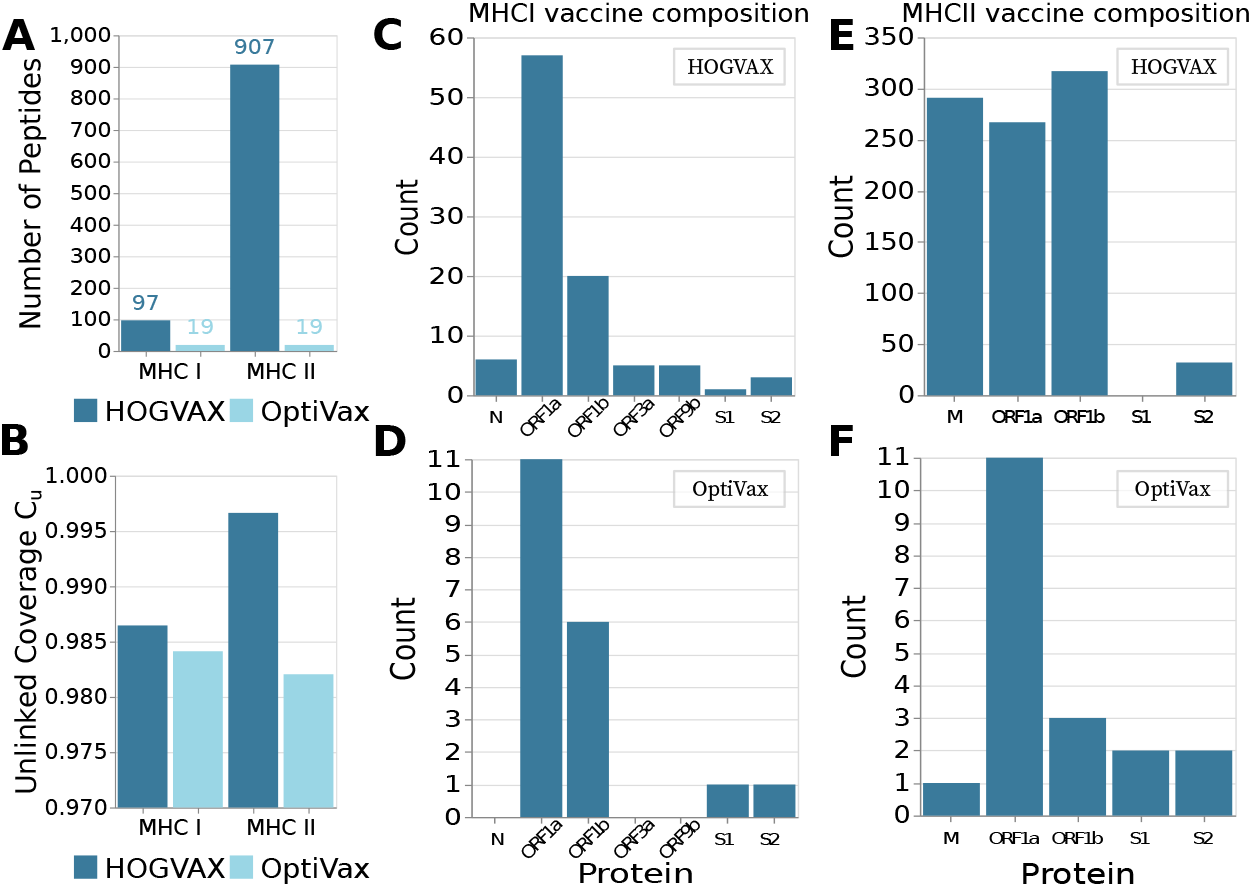
Comparison of HOGVAX and OptiVax-Unlinked vaccines on single-allele data. **A** Number of peptides included in the vaccine candidates. **B** Unlinked population coverage *C*_*u*_ of the average world population predicted with EvalVax-Unlinked. **C, D** Peptide viral protein origins of the MHC class I vaccines. **E**,**F** Peptide viral protein origins of the MHC class II vaccines.

With EvalVax-Unlinked, we achieved a population coverage of 98.41 % for the MHC class I vaccine set of OptiVax-Unlinked and slightly higher coverage of 98.65 % for the HOGVAX vaccine, see Fig. 3B.

For the MHC class II vaccine, OptiVax-Unlinked created a set of 19 peptides of total length 322. Thus, we called HOGVAX with *k* = 322 and obtained a vaccine set of 907 peptides. With EvalVax-Unlinked, we predicted a population coverage of 98.2 % for the OptiVax-Unlinked-designed vaccine, and a slightly higher coverage of 99.66 % for the HOGVAX-designed vaccine of MHC class II, see Fig. 3B. Although the amount of peptides in our vaccine candidate is almost 50 times greater than that of OptiVax-Unlinked, fewer different proteins are represented in the vaccine of HOGVAX. More precisely, HOGVAX renounced peptides of the S1 unit of the spike protein and the distribution of peptides from distinct proteins is also different from the distribution in the OptiVax-Unlinked vaccine. For example, the second largest fraction of peptides in the HOGVAX vaccine derives from the M protein, where the second largest number of peptides chosen by OptiVax-Unlinked originates from ORF1b. The exact compositions are given in Fig. 3C.

We also evaluated the robustness of the computed peptide sets using the EvalVax-Robust metric on individual haplotype data. The probabilities of each number of observed peptide-HLA hits of the MHC I vaccines for HOGVAX and OptiVax-Unlinked are presented in Fig. 4A and B, respectively. HOGVAX achieved an overall higher number of expected peptide-HLA hits in each of the three populations of Europeans, Africans, and Asians. Moreover, the maximum value of observed hits is higher by approximately ten hits compared to the OptiVax-Unlinked outcome. This is also evident in the predictions calculated with EvalVax-Robust that are presented in Fig. 4C and D for HOGVAX and OptiVax-Unlinked, respectively. With the HOGVAX-designed vaccine, we obtained 99.998 % coverage for at least one peptide hit, 93.243 % for at least five hits, and 47.676 % for at least ten hits for the average of all three populations. In contrast, we observed a stronger decrease in the coverage achieved by the OptiVax-Unlinked vaccine candidate for higher hit counts. For the average population, the OptiVax-Unlinked peptides were predicted with 99.994 % coverage for at least one peptide-HLA hit, 87.538 % coverage for at least five hits, and 20.377 % coverage for at least ten hits. The improvements are even more pronounced for the MHC II data set where HOGVAX achieved an expected number of 176.41, 176.76, and 101.9 peptide-HLA hits per individual of the European, African, and Asian population, respectively. The OptiVax-vaccine candidate achieved 9.78, 9.67, and 6.6 per-individual peptide-HLA hits for Europeans, Africans, and Asians, respectively. Accordingly, the probability of at least ten peptide hits per individual in an average population was higher for the HOGVAX vaccine with 99.457 % compared to 38.639 % robust coverage for the OptiVax vaccine.

**Fig. 4.**
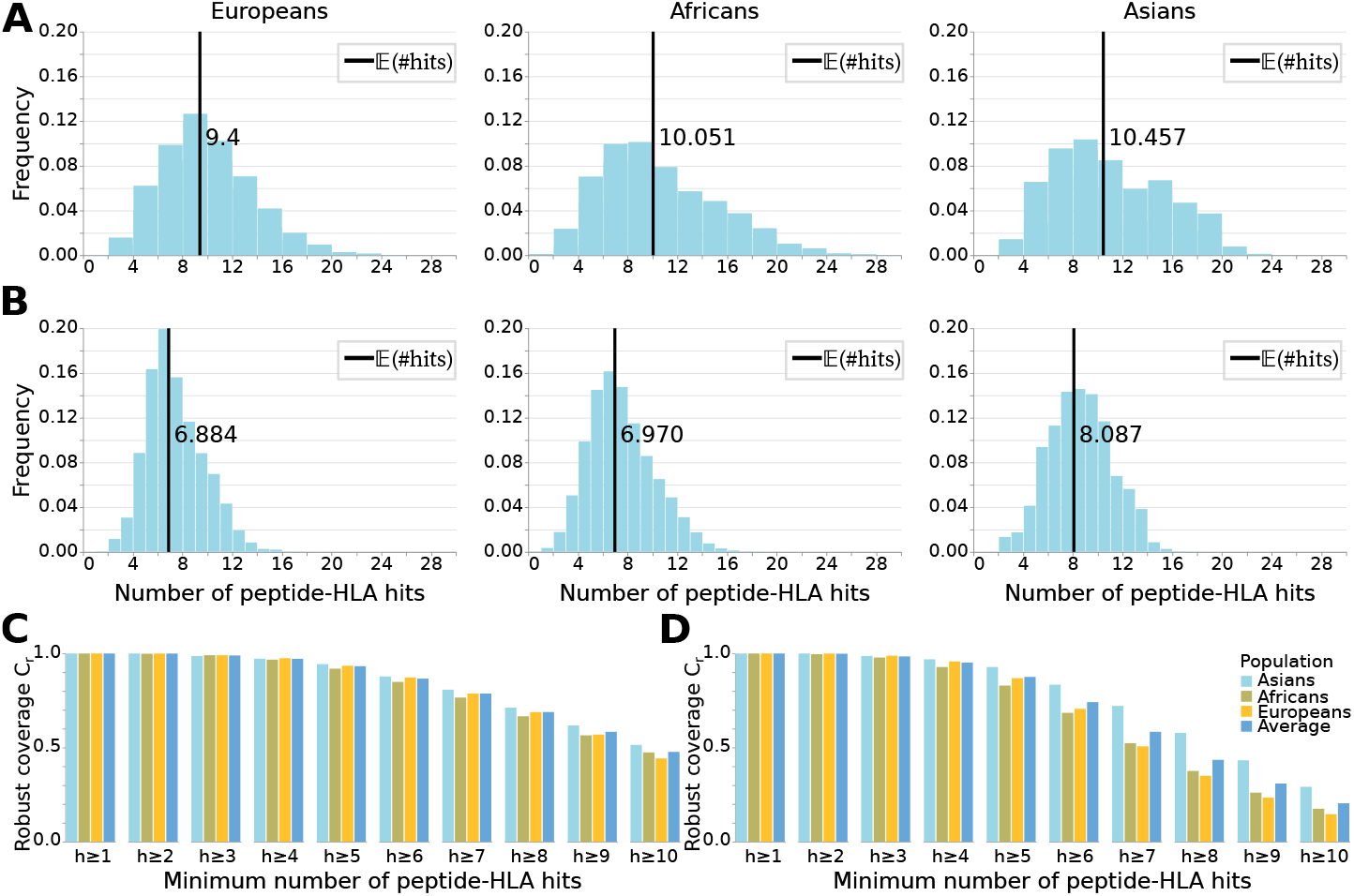
Evaluation of population coverage of the MHC class I vaccine candidates optimized on single-allele data for populations self-reporting as having European, African, or Asian ancestry. **A** HOGVAX and **B** OptiVax-Unlinked distribution of the number of peptide-HLA hits and expected number of hits per individual for the three populations. **C** HOGVAX and **D** OptiVax-Unlinked population coverage computed with EvalVax-Robust for the three populations and average values.

### 4.6 Vaccine design based on haplotype frequencies

Here, we used haplotype frequencies of three populations self-reporting as having European, African, or Asian ancestry. To determine the maximum length of our vaccine candidate, we ran OptiVax-Robust on haplotype frequencies resulting in a set of 19 peptides. From their concatenation, we inferred a maximum vaccine length of *k* = 174 for MHC class I. Within this limitation, HOGVAX selected 123 peptides for which we predicted an unlinked coverage of 97.56 %. Using EvalVax-Robust, we calculated a robust coverage of 99.998 %, 86.844 %, and 32.694 % for at least one, five, and ten peptide-HLA hits per individual of an average population.

Following the same approach, we determined a maximum length of *k* = 324 for the HOGVAX MHC class II vaccine sequence. HOGVAX chose 414 peptides that achieved an unlinked coverage of 99.49 %. We calculated expected per-individual hit numbers of 63.1, 68.1, and 28.93 for the European, African, and Asian population. Accordingly, the robust coverage for an average world population is significantly higher with 96.667 % for at least ten peptide-HLA hits per individual.

## 5 Discussion and Conclusions

We present HOGVAX, a novel combinatorial approach for MHC class I and II peptide vaccine design that solves the *Maximum Scoring k-Superstring* problem. As the underlying data structure we use the hierarchical overlap graph. With HOGVAX, we intend to fill the gap of uncovered rare alleles, maximize population coverage based on allele frequencies of the target population, and induce robust immunity against virus variants by many per-individual peptide-HLA hits. Due to our overlap vaccine design, we can fit a significantly larger amount of peptides within the size of a regular vaccine constructed from concatenated peptides. In this work, we compare HOGVAX to OptiVax by Liu et al. [13]. Even though the MSKS problem is NP-hard, we are able to create provably optimal vaccine candidates in reasonable run time for realistic instance sizes using our HOG-based ILP. In contrast, OptiVax is based on a heuristic search that does not necessarily return an optimal peptide vaccine set.

Note that OptiVax chooses a specific number of peptides and Liu et al. did not assume any cutting of the peptides. Our sequence, however, must be cut into peptides. We are therefore able to select more peptides than in the OptiVax vaccine. As reflected by our results, the larger number of peptides chosen by HOGVAX generally leads to higher predicted population coverage and per-individual peptide-HLA hits compared to the OptiVax-designed vaccines. To not waste any space, our model likely chooses peptides that cover multiple loci or haplotypes at once. This, and the high number of peptides would explain the tremendous increase of expected hits. We assume that a higher number of peptide-HLA hits per genotype correlates with higher efficacy of the vaccine [13], for example by inducing immunity against various parts of the pathogen. Moreover, a large set of peptides leads to a more conserved immunity to viral mutations, as the probability of simultaneous mutations in the viral peptides exponentially decreases with an increasing number of considered peptides [21].

The diverse compositions of the HOGVAX and OptiVax vaccines are worth mentioning. HOGVAX often chooses more peptides from the ORF1a and the membrane protein compared to OptiVax. Grifoni et al. showed that the largest proportion of peptides recognized by CD4^+^ and CD8^+^ T cells in individuals unexposed to SARS-CoV-2 derived from ORF1a and ORF1b [10]. The second largest fraction of recognized peptides originated from the spike protein, followed by peptides from the membrane protein [10]. However, T cells from individuals infected with COVID-19 mostly targeted the spike protein, followed by similar fractions of the peptides from the membrane protein and ORF1a and b [10]. It seems like the vaccine compositions model the T cell activation of unexposed individuals. From the SARS-CoV outbreak in 2002, we know that we need to vaccinate against the spike protein to prevent infections [6]. However, the amount of S1 and S2 peptides in the vaccine candidates computed with HOGVAX and OptiVax is rather low, which is likely the cause of the specific predictions of NetMHCpan-4.1 and NetMHCIIpan-4.0. Nevertheless, by targeting further proteins, a vaccine can be more robust against viral mutations that otherwise likely lead to immune escapes as in the case of the SARS-CoV-2 Omicron variant and the available spike protein vaccines [22].

We realized that multiple optimal HOGVAX solutions exist. Exploring and structuring the space of optimal solutions is part of our future work, including the investigation and elimination of symmetric solutions that correspond to traversing the same closed walk in different orders. Multiple solutions also offer new opportunities to further improve our method. For example, we could maximize the number of peptides in the output set in a second optimization, where we fix the already optimized population coverage by a constraint. Another possibility would be to also optimize the cleavage probability of peptides. We could also include peptides that were shown to be good vaccine candidates by fixing the corresponding variables to build a vaccine sequence around the pre-selected peptides. Furthermore, HOGVAX is not restricted to the application to SARS-CoV-2, but could, for example, be used to design cancer vaccines. Peptide vaccines are considered a promising approach in oncology for cancer prevention and therapy [14].

In this work, we used single allele and haplotype frequencies to optimize our peptide vaccines. In the future, we want to work on the scalability of HOGVAX, for example, towards diplotype frequency data which can be constructed from the Cartesian product of the haplotypes for each HLA class independently.

So far, however, little is known about the protein degradation by the proteasome, such that there is uncertainty if the peptides of our overlap-vaccine sequence will be present in the end. Thus, only experimental investigations will show how well our approach and competing methods will work in practice.

To our knowledge, HOGVAX is the first method that designs peptide vaccines by exploiting overlaps between the peptide sequences. Thereby, we significantly increase the number of peptides included in a vaccine and suggest a strategy to potentially conquer the challenge of high HLA diversity. With our vaccine formulations, we maximize population coverage and induce robust immunity against SARS-CoV-2. Our approach is flexible and can easily be adjusted for other purposes, like vaccine design against novel SARS-CoV-2 variants and other diseases like cancer.

